# Near-surface flight enhances thrust in an insect-inspired flapping-wing robot

**DOI:** 10.64898/2026.11.24.768668

**Authors:** Chenyu Zhang, Cameron Urban, Hang Gao, Z. Jane Wang, E. Farrell Helbling

## Abstract

A number of semi-aquatic insects rapidly move across the water surface using interfacial flight. In this locomotion strategy, flapping wings generate thrust above the air-water interface while surface tension supports their weight. This close proximity to the air-water interface alters aerodynamic force generation through ground effect, potentially enhancing horizontal propulsion. Here, we demonstrate that near-surface flapping enhances horizontal thrust by up to 17% at shallow stroke-plane inclinations. Using a beetle-inspired flapping-wing robot, we systematically vary wing-surface distance (*D*/*c*, where *D* is the minimum trailing-edge-to-surface clearance during flapping, normalised by chord length *c*) and stroke plane inclination (*β*) at Reynolds numbers of order 10^3^. Locomotion experiments show increased acceleration and higher terminal velocity near the surface (*D*/*c* < 1) at shallow stroke plane angles (*β* = 30° and 45°), while direct wind-tunnel force measurements confirm increased aerodynamic thrust, distinguishing this enhancement from changes in water-related resistance. Particle image velocimetry further reveals that near-surface flapping alters passive wing motion and surrounding flow, producing stronger backward-directed flow after stroke reversal and earlier leading-edge vortex development during the initial upstroke. These findings establish a configuration-dependent aerodynamic benefit of near-surface flapping, providing insight into insect-scale interfacial flight and opportunities for enhancing propulsion in bio-inspired robotic systems.

## 1. Introduction

Animals frequently fly close to ground or water during behaviours such as drinking, foraging, and takeoff [1, 2, 3, 4]. In addition to the behavioural advantages of operating near a surface, aerodynamic ground effect can modify force generation and reduce the energetic demands of flight [5, 6]. At the insect scale, waterlily beetles (*Galerucella nymphaeae*) exhibit interfacial flight, moving rapidly along the air-water interface using flapping wings while their legs remain in contact with the water and their body weight is supported by surface tension [7]. Despite the additional hydrodynamic resistance associated with water contact, these beetles frequently employ interfacial flight. This behaviour raises the question of whether flapping near the water surface provides an aerodynamic advantage through enhanced horizontal thrust, potentially complementing other benefits of remaining at the interface.

Ground effect has been extensively studied in engineered aerodynamic systems, from fixed wings to oscillating foils and flapping-wing vehicles. In fixed-wing flight, proximity to a boundary can increase lift and reduce drag [8]. In unsteady propulsion, nearby boundaries interact with vortices generated by oscillating wings, modifying aerodynamic force production. Studies of heaving and pitching foils have demonstrated that these interactions can enhance thrust and alter lift [9, 10, 11]. More closely resembling insect flight, root-pivoted flapping wings have been analyzed during **2** takeoff [12, 13, 14], hovering [15], and forward flight [16]. These studies demonstrate that ground proximity can modify aerodynamic forces and vortex development, with the response affected by boundary clearance, wing kinematics, and Strouhal number (the ratio of flapping velocity to forward speed).

During interfacial flight, insect wings operate at shallow stroke-plane angles (*β* < 45°) and undergo passive wing rotation at Reynolds numbers of order 10^3^ [7]. Stroke-plane inclination changes the distribution of wing-surface clearance along the wingspan and throughout the flapping cycle. At shallow inclinations, the wing can remain close to the surface over a longer portion of the stroke, potentially increasing aerodynamic interactions with the boundary. Previous studies have examined inclined stroke planes during near-surface hovering [17, 18, 19], while previous studies of forward propulsion have often employed prescribed pitching and heaving motions or fixed stroke plane configurations [10, 11, 16, 20]. Furthermore, passive wing rotation allows aerodynamic loading to modify wing orientation and subsequent force generation, introducing additional coupling between wing motion and surrounding flow [21, 22]. How wing-surface proximity and stroke-plane inclination affect horizontal thrust, passive wing motion, and near-surface flow during insect-scale forward flight remains unresolved, motivating a systematic experimental characterization of these interactions. Directly characterizing ground effect in living insects is challenging because wing-surface distance, stroke-plane inclination, and wing kinematics vary during locomotion and are difficult to control independently. Measuring the small aerodynamic forces generated by insects presents an additional challenge [23]. Robotic physical models offer an alternative, enabling controlled and repeatable studies of flapping-wing aerodynamics [24, 25, 26] and insect-inspired manoeuvres [27, 28, 29]. Here, we use a sub-gram beetle-inspired robot [30] with flapping wings for propulsion and hydrophobic legs to support the robot on the water surface (figure 1*a*). We can independently adjust the wing-surface distance and stroke-plane inclination while retaining passive wing rotation. We combine three complementary approaches to characterize the effects of ground proximity on horizontal propulsion: straight-line locomotion experiments with dynamic modelling, direct wind-tunnel thrust measurements under controlled relative airflow, and particle image velocimetry (PIV) to examine wing kinematics and surrounding flow.

**Figure 1.**
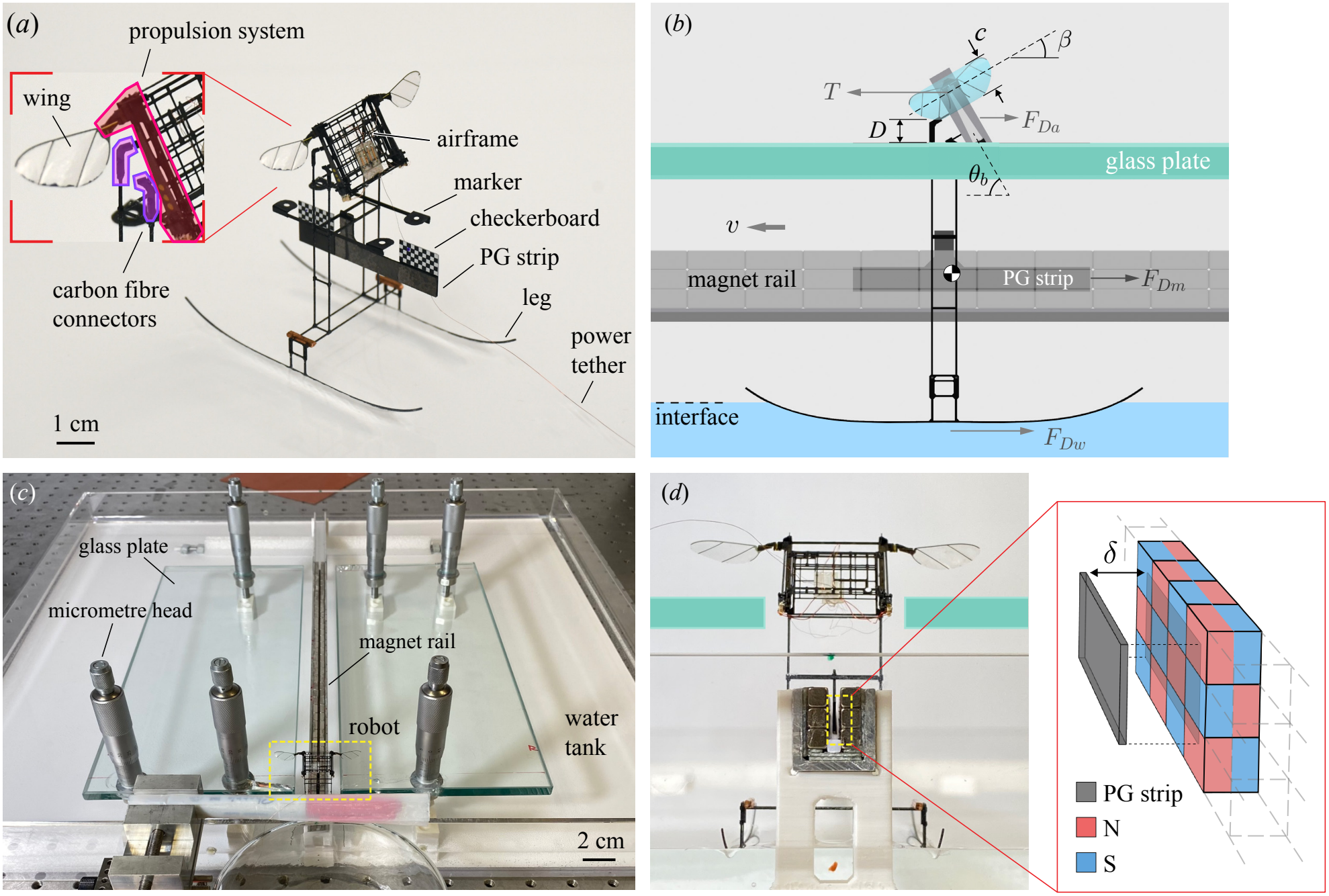
Beetle-inspired flapping-wing robot and straight-line locomotion apparatus. (*a*) The beetle-inspired robot resting on the water surface. The propulsion system consists of the wing, actuator, and transmission. Carbon fibre connectors on the airframe vary body pitch angle. The hydrophobic legs maintain contact with the water surface, and a power tether supplies actuation signals. The markers facilitate motion tracking and the checkerboard provides a visual reference for body pitch and height. (*b*) Side-view schematic showing the minimum trailing-edge clearance *D*, stroke-plane inclination *β*, body pitch angle *θ*_*b*_, and horizontal forces acting on the robot during forward motion including thrust *T*, aerodynamic drag *F*_*Da*_, hydrodynamic drag *F*_*Dw*_, and magnetic drag *F*_*Dm*_. The robot moves with a forward velocity *v*. (*c*) Experimental apparatus with adjustable-height glass plates beneath the wings and a noncontact magnetic guide rail for straight-line locomotion. The robot is highlighted in the yellow box for scale. (*d*) Cross-sectional schematic of the magnetic guide rail. Diamagnetic repulsion centres the PG strip between two permanent magnet arrays (alignment highlighted in the inset), constraining lateral motion at a fixed gap *δ* without mechanical contact (see **electronic supplementary material, movie S1**).

Using this experimental framework, we quantify the aerodynamic benefits of ground proximity during insect-scale forward flight. At a shallow stroke-plane inclination (*β* = 30°), reducing the normalized wing–surface distance from *D*/*c* =9 to 0.2 increases horizontal thrust by approximately 17%. This enhancement is strongly dependent on stroke-plane inclination, highlighting the importance of wing configuration in determining the aerodynamic response to nearby boundaries. Our findings provide experimental evidence that ground effect can enhance horizontal propulsion during insect-scale flapping and suggest a potential aerodynamic advantage for insects performing interfacial flight.

## 2. Methods

We use an insect-scale flapping-wing robot as a physical model to study aerodynamic ground effect during interfacial flight. We develop a straight-line locomotion apparatus that allows independent adjustment of wing–surface distance and stroke-plane inclination, and formulate a longitudinal dynamic model to estimate thrust from robot motion. We also construct a wind-tunnel apparatus with controlled airflow and moving boundaries for direct thrust measurements. Finally, we use particle image velocimetry (PIV) to measure near-wing flow and reconstruct sectional wing kinematics.

### 2.1. Robotic platform

The beetle-inspired robotic platform is comprised of an airframe, hydrophobic legs to support the weight of the robot on the water surface, and a propulsion system (adapted from prior work [30]). We fabricate these components using the Smart Composite Microstructures process, described in detail in [31]. We complete all laser patterning steps using the Protolaser U4 Laser cutter (LPKF, 20 µm beam width), and apply the necessary temperature and pressure profiles for laminate manufacturing with a Carver press (Model 4386). The airframe uses a 165 µm, three-ply carbon fiber laminate (M55J, TorayCA) orthogonally stacked to provide high bending stiffness against actuator loads and external disturbances. The frame connects the two flapping wing propulsion systems and the legs (see Fig. 1(a)). We fabricate the legs using a five-ply carbon fiber laminate (M55J) orthogonally stacked to increase in-plane stiffness along the forward skating direction. Patterned legs are coated in hydrophobic coating (Ultra-Ever Dry) for a final leg thickness of 0.324 mm. The legs can support up to 1.5 g of payload using surface tension, well beyond the total mass of the system (759 mg). We hand-assemble and connect all components with cyanoacrylate glue (Loctite 416, 495) unless otherwise noted. We supply power to the robot using a four-wire bundle of magnet wire (52-gauge; part number 52TPN-155 Red, MWS), selected to minimize torque on the robot during operation.

Each flapping-wing propulsion system consists of a piezoelectric bimorph actuator (design and fabrication detailed in [32]), transmission, passive wing hinge, and wing. We fabricate the propulsion system using the process detailed in [33] and select the transmission stiffness and wing hinge stiffness to achieve the desired wing stroke motion. Each wing has a length of *R* = 15 mm and a maximum chord *c* = 5 mm. The wing hinges permit passive rotation about the spanwise axis (wing pitch), allowing aerodynamic and inertial loading to determine wing pitch during each flapping cycle. We operate the system at its mechanical resonance with a wing stroke peak-to-peak amplitude of 44°, wing pitch angle of 62° (at mid-stroke), and frequency of 170 Hz. Importantly, we observed no change in wing kinematics between the start and end of the experiments.

To vary stroke-plane inclination, we used customized carbon-fibre connectors to set the body pitch angle *θ*_*b*_ (figure 1*c*). For our robot configuration, the stroke-plane inclination satisfies *β* = 90°*−θ*_*b*_. We varied *θ*_*b*_ from 0° to 60° in 15° increments, corresponding to *β* = 90° to 30°. The shallow configurations (*β* = 30° and 45°) approximate the stroke-plane inclinations observed in waterlily beetles during interfacial flight [7], while the steeper configurations provide comparisons across a wider range of wing orientations.

### 2.2. Straight-line locomotion experiments

To systematically assess the effects of wing-surface distance and stroke-plane inclination on robot locomotion, we developed a straight-line testing apparatus with adjustable rigid boundaries and noncontact magnetic guidance (see figure 1*b-d*)). Two glass plates positioned beneath the wings provide rigid boundaries that isolate aerodynamic ground effect from deformation of the water surface. Each plate is supported by three micrometre heads with a positioning accuracy of 10 µm, allowing precise adjustment of wing-surface clearance while maintaining the robot configuration.

To constrain the robot to one-dimensional motion and compensate for potential manufacturing asymmetries between the wings, we incorporated a noncontact magnetic guide rail. A pyrolytic graphite (PG) strip (38 mm *×* 4.5 mm *×* 0.4 mm), mounted between the robot legs, moves within a channel formed by two arrays of permanent magnets. The arrays face each other with opposing polarities, generating diamagnetic repulsion that passively centres the strip within the channel [34, 35]. The strip maintains a 0.5 mm lateral clearance and remains clear of the channel bottom, as verified by high-speed videography (see **electronic supplementary material, movie S1**).

We define the wing-surface distance *D* as the minimum trailing-edge clearance at the lowest point of the wing stroke (figure 1*c*), measured from high-speed images acquired with a macro lens (VEO 710L, Phantom; Irix USA). We normalize the distance by the local wing chord *c* at the radius of the second moment of wing area *r* = 0.6*R*, where *r* the spanwise location measured from wing root. This definition allows direct measurement of clearance without requiring localization of the wing hinge. We recorded robot displacement using the high-speed videography (frame rate of 1000 fps) and aligned the focal plane with the tracking markers attached to the robot (figure 1*a*) and parallel to the water surface. Given a camera resolution of 832 *×* 800 pixels, we achieved a spatial resolution of 0.6 mm per pixel after distortion correction and spatial calibration. We extracted robot displacement using custom software that leverages the “image processing toolbox” in MATLAB (R2024a, MathWorks). The process is as follows: we manually select a small region around each marker in the first video frame; the program then automatically updates each region’s location to follow the moving marker. In each frame, the program thresholds the image to isolate the markers, locate the centroids of each marker, and calculate the displacement from previous frames.

We tested five stroke-plane angles from *β* = 30° to 90° in increments of 15°, with five normalised wing-surface distances (*D*/*c*) for each angle, spanning near- and far-surface conditions up to *D*/*c ≈* 4. For each condition, we conducted three trials in which the robot accelerated from rest and travelled approximately 40 cm, reaching terminal velocity before reaching a physical stop. We calculated forward velocity (*v*) by applying central differences to the measured displacement and then filtering the velocity using a second-order Butterworth low-pass filter with a cutoff frequency of 15 Hz. We report the mean and standard deviation of the velocity trajectories across the three trials.

### 2.3. Wind-tunnel apparatus

To measure aerodynamic thrust and visualize the near-wing flow pattern under controlled forward relative motion, we developed a wind-tunnel apparatus with adjustable wing-surface clearance and moving ground boundaries (figure 2). The blowdown wind tunnel consists of a bell-mouth inlet, a PWM-controlled fan (P14 Slim, ARCTIC), a flow-conditioning section, a contraction section, and a test section with a cross-sectional area of approximately 95 mm *×* 95 mm. We included a honeycomb as well as coarse and fine screens in the flow-conditioning section to reduce turbulence, while the contraction improves flow uniformity. We controlled the freestream velocity by varying the pulse-width-modulated duty cycle. We calibrated the velocity using high-speed videography of generated glycerin-based smoke streaks [36]. We can adjust the freestream velocity between 0.12 m s^−1^ and 1.00 m s^−1^ in approximate increments of 0.01 m s^−1^, with less than 10% spatial variation across the central 80% of the test-section cross-section.

**Figure 2.**
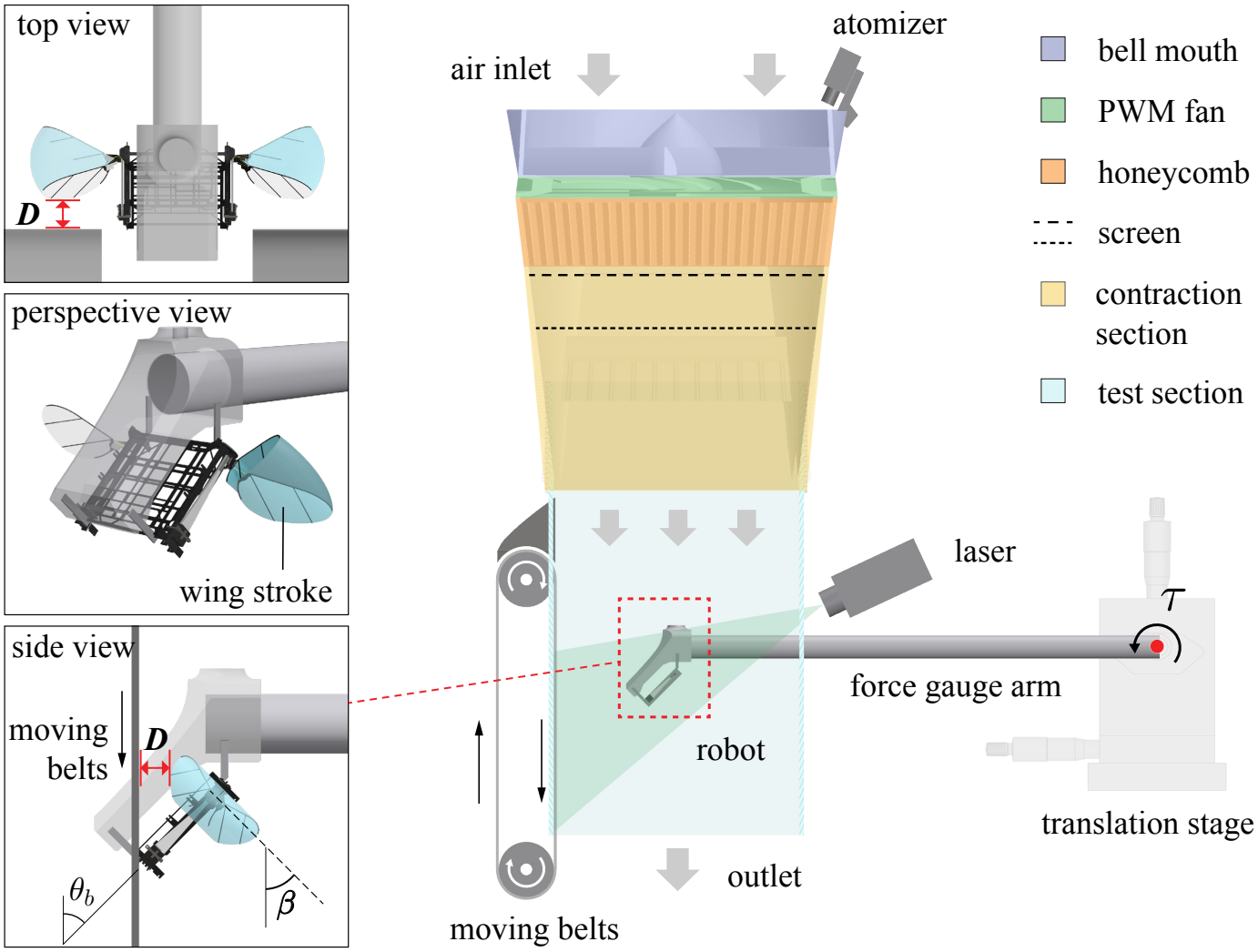
Illustration of the wind-tunnel apparatus for thrust measurements and particle image velocimetry (PIV). The blowdown wind tunnel provides controlled airflow over the flapping-wing robot, which is mounted on a lever arm connected to an external force/torque sensor. Custom mounting fixtures set the body pitch angle *θ*_*b*_, while a translation stage controls the wing-surface distance *D* from the moving belts. These belts match the freestream velocity to reproduce the relative motion between the robot and underlying surface during forward locomotion. An atomizer seeds the airflow with tracer particles illuminated by a laser sheet for PIV measurements. Insets illustrate the robot mounting geometry and wing stroke range (blue); passive wing rotation is not shown. Schematics are not to scale. Set-up and operating demonstrations are detailed in **electronic supplementary material, movie S2**.

We mounted the robot body at one end of a 22.5 cm carbon-fibre arm that extended into the test section, with the other end attached to an external force/torque sensor (Nano17 Ti, ATI, resolution 0.1 mN). The arm transmits the aerodynamic thrust generated by the flapping wings as a torque measured by the sensor. Custom mounting fixtures set the robot orientation to match the stroke-plane angles used in the straight-line locomotion experiments. A three-axis translation stage adjusted the robot position in the section and wing-surface clearance. To minimize wall effects of the test section, we centred the robot with wall clearances exceeding 8*c*.

To reproduce the relative motion between the robot and the surface during forward locomotion, we installed two moving belts [37] beneath the wings. We matched the belt speed (controlled with a PWM motor) to the freestream velocity, reducing the development of a stationary-wall boundary layer that could influence near-surface wing aerodynamics. We measured wing-surface distance *D* relative to the belt surface using the definition in §2.2. This arrangement enables direct thrust measurements and particle image velocimetry under controlled airflow velocity, stroke-plane inclination, and boundary clearance.

### 2.4. Aerodynamic thrust measurements

We measured the mean aerodynamic thrust generated by the robot using the lever-arm force/torque sensor described in the previous section (§2.3). We calibrated the sensor by applying known static loads to the lever arm, covering the range of expected force measurements, 0.3–2.7 mN.

For each condition, we averaged the thrust over a 0.8 s interval (~130 cycles) of steady-state flapping. The limited dynamic bandwidth of the lever-arm system prevented us from reliably reconstructing instantaneous forces within individual wingbeats. We repeated each measurement twice, with less than 3% variation between the resulting mean thrust values.

### 2.5. Longitudinal dynamic model

To estimate the thrust from the robot position in straight-line experiments, we model the robot forward dynamics [30] along the magnetic guide rail by:

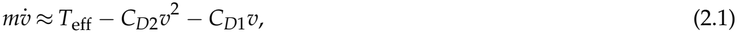

where *m* is the robot mass, *v* is the forward velocity, and *T*_eff_ is the effective aerodynamic thrust, which depends on wing–surface clearance, stroke-plane inclination, and forward velocity. The quadratic term represents the combined hydrodynamic and aerodynamic resistance (*F*_*Dw*_ and *F*_*Da*_, respectively), while the linear term accounts for eddy-current damping (*F*_*Dm*_) introduced by the magnetic guide rail [38] (see figure 1*b*). Because the quadratic resistance is dominated by water drag [30], we assume that the resistance coefficients remain independent of wing-surface distance, as the water contact geometry remains constant during the tests.

To identify the resistance coefficients, we first measured the baseline robot thrust *T*_0_(*v, β*) under far-from-surface conditions (*D*/*c* > 8) using the wind tunnel apparatus described in the previous section. At this distance, we assume *T*_eff_ *≈ T*_0_(*v, β*). For each angle *β*, we fit the corresponding thrust curves *T*_0_(*v*) using a third-order polynomial (figure 3*a*). This fitting is consistent with the theoretical dependence of mean thrust on forward velocity at small advance ratios (*J* = *v*/*w* < 1) and fixed cycle-averaged wing sweep speeds (*w*; see **electronic supplementary material** for detailed justification). We then conducted straight-line control experiments at *β* = 30° without the magnetic guide rail to determine the quadratic resistance term *C*_*D*2_ due to hydrodynamic and aerodynamic drag. We then reintroduced the magnetic rail to determine the additional linear resistance coefficient *C*_*D*1_, keeping *C*_*D*2_ constant (there is no observable change in leg meniscus height, see **electronic supplementary material, movie S3**). We identified both coefficients using nonlinear grey-box identification in MATLAB, numerically integrating the longitudinal model (equation (2.1)) and minimizing the squared differences between measured and fit velocity trajectories (figure 3*b,c*). We obtained *C*_*D*2_ = 4.815 g m^−1^ and *C*_*D*1_ = 0.520 mN s m^−1^.

**Figure 3.**
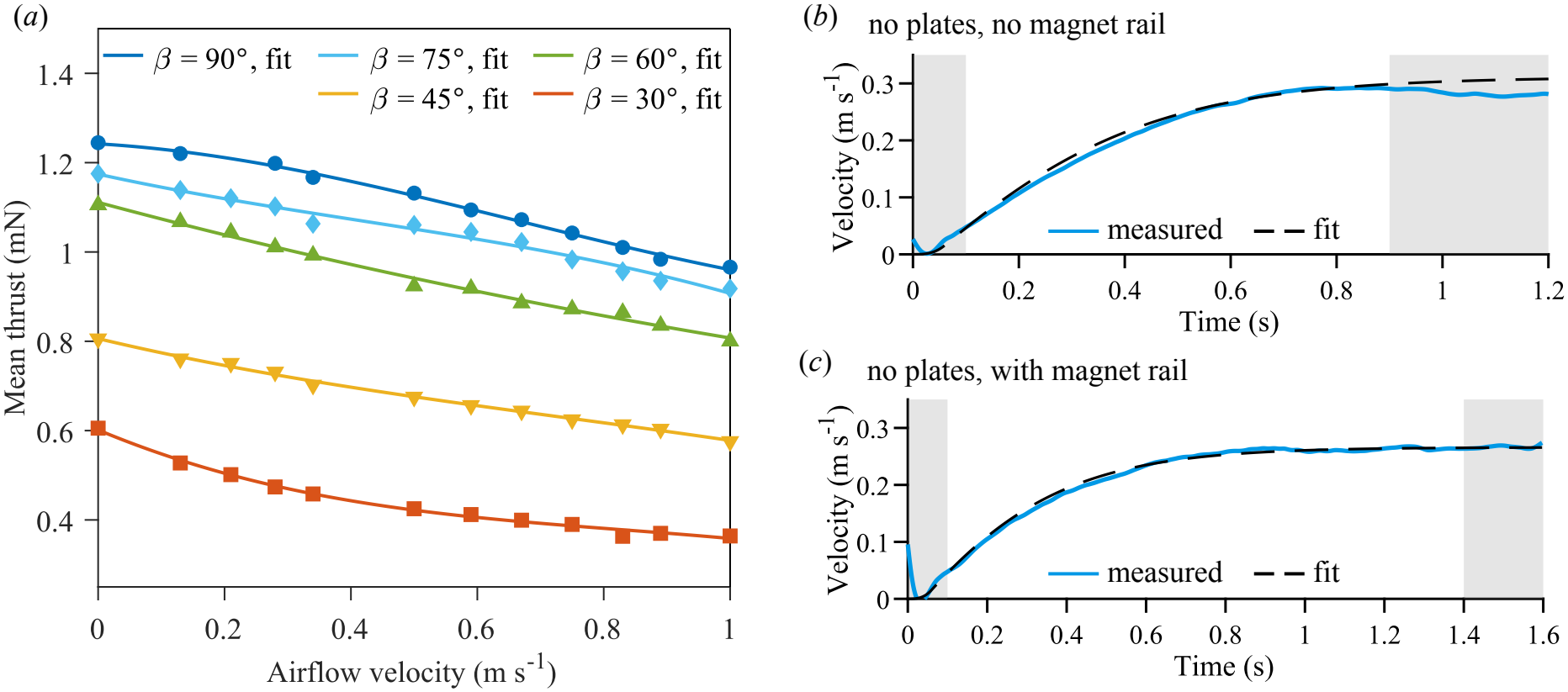
Baseline thrust characterization and longitudinal model calibration. (*a*) Mean aerodynamic thrust as a function of freestream velocity for different stroke-plane angles *β* under far-from-surface conditions (*D*/*c* > 8). Mean computed over two trials, with approximately 130 flapping cycles per trial. We define the baseline thrust-velocity relationship *T*_0_(*v*, *β*) using a third-order polynomial fit (solid curves). (*b*) Identification of the quadratic drag coefficient (*C*_*D*2_) at *β* = 30*°*, far-from-surface, and rail-free condition. (*c*) Identification of the rail-induced linear resistance coefficient (*C*_*D*1_) with the rail installed. In (*b,c*) we present the fitted velocity trajectories with the calibrated longitudinal model (dashed line), compared with measured trajectories (blue line). Grey-shaded regions indicate time intervals excluded from parameter identification due to initial filtering transients at the start of the trial and contact between the wire tether and the water at the end of a trial, which introduces an external torque not captured by the model.

We focused the model-based thrust estimation on *β* = 30°, which exhibited the strongest locomotor response to wing–surface proximity and approximates the stroke-plane inclination observed in waterlily beetles during interfacial flight [7]. We estimated the effective aerodynamic thrust *T*_eff_ from the measured terminal velocity *v*_ter_, where the longitudinal acceleration approaches zero. Under this condition, the force balance reduces to:

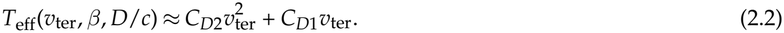

We compared this estimate with the baseline thrust *T*_0_(*v*_ter_, *β*) at the terminal velocity to assess the effect of wing-surface proximity.

### 2.6. Particle image velocimetry

We performed two-dimensional PIV to characterize the flow around the flapping wings (figure 4*a*). A 520 nm, 1 W continuous-wave laser generated a light sheet less than 1 mm thick, intersecting the wing at a nominal location of *r* = 0.6*R* (figure 4*b*). We seeded the airflow with droplets (diameter *≈* 5 µm) using piezoelectric atomizers (SJ56, SazkJere).

**Figure 4.**
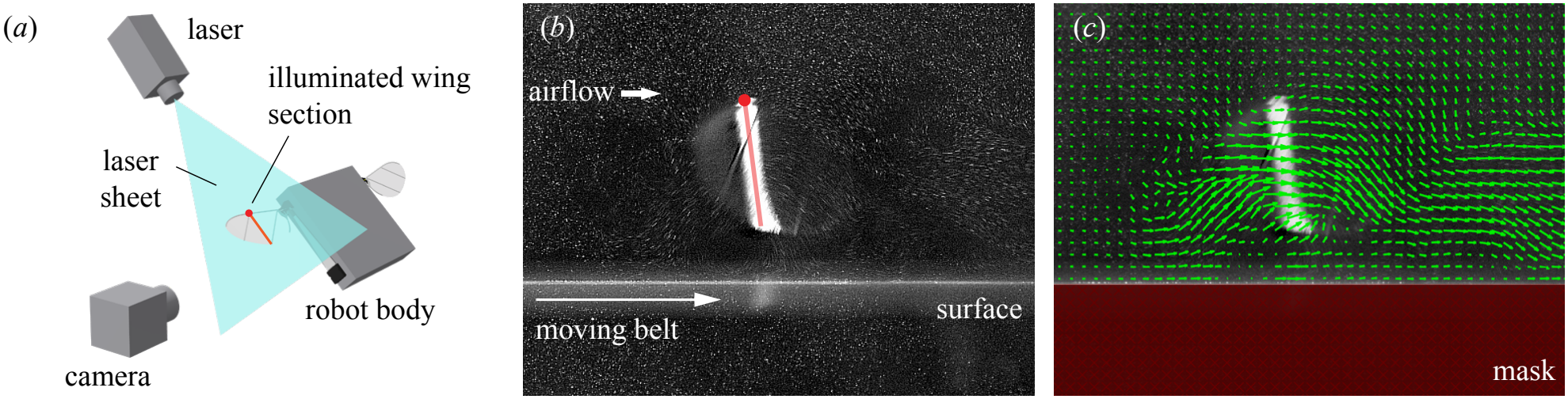
PIV configuration and processing. (*a*) Schematic of the PIV setup, with a laser sheet intersecting the flapping wing at *r* = 0.6*R*. (*b*) Representative high-speed image of the illuminated wing section and seeded airflow. The red line indicates the detected wing section, and the dot marks the leading edge. (*c*) Corresponding velocity field (green arrows) reconstructed from digital PIV analysis, with masked regions outside the area of interest. (*b*–*c*) show the flapping wing shortly after the downstroke–upstroke reversal.

Using high-speed videography, we recorded particle images at 11 900 fps, corresponding to 70 frames per wingbeat.

A real-time control system (Baseline S, Speedgoat) synchronized the camera trigger and actuation signals to provide a consistent timing reference across experiments. We achieve a spatial resolution of 33 µm per pixel.

We processed the images in PIVlab (version 3.12, MATLAB) [39]. We first applied contrast-limited adaptive histogram equalization (CLAHE) to improve particle visibility, then calculated velocity vectors using a multipass Fast Fourier Transform with an interrogation window of 24 *×* 24 pixels and 50% overlap. We used the resulting velocity fields (figure 4*c*) to compare near-wing flow structures and estimate momentum-flux. The vorticity field is obtained by spatial differentiation of the velocity field.

Finally, we reconstructed sectional wing kinematics from the PIV images. As the wing sweeps through the fixed laser sheet, the illuminated section can shift along the span during the stroke. Keeping the plane fixed and tracking the illuminated intersection can be used as a reasonable approximation for the wing’s position and orientation throughout the flapping cycle [22, 25].

## 3. Results

We demonstrate that near-surface flapping enhances horizontal propulsion in an insect-scale robot, with the strongest effects at shallow stroke-plane angles. Straight-line locomotion experiments reveal increased acceleration and terminal velocity, while model-based thrust estimates and direct wind-tunnel measurements provide complementary evidence of enhanced aerodynamic thrust. We then examine changes in passive wing kinematics and near-wing flow to identify aerodynamic features associated with this enhancement. Finally, momentum-balance estimates from PIV provide a quantitative link between the altered flow and thrust generation.

### 3.1. Ground effect on straight-line motion

Wing-surface proximity increased robot acceleration and terminal velocity during straight-line locomotion, with the strongest effects at shallow stroke-plane angles, as shown in figure 5. At *β* = 30° and 45°, the robot accelerated more rapidly and reached higher terminal velocities as *D*/*c* decreased (figure 5*a-b*). The enhancement was most pronounced at *D*/*c* < 0.7. In contrast, the velocity responses at *β* = 90° remained consistent across the tested distances (figure 5*c*), while intermediate angles (*β* = 60° and 75°) exhibited similarly weak dependence on surface proximity. The higher terminal velocity at *β* = 90° is expected given the larger horizontal component of aerodynamic force, which provides greater thrust (figure 3*a*).

**Figure 5.**
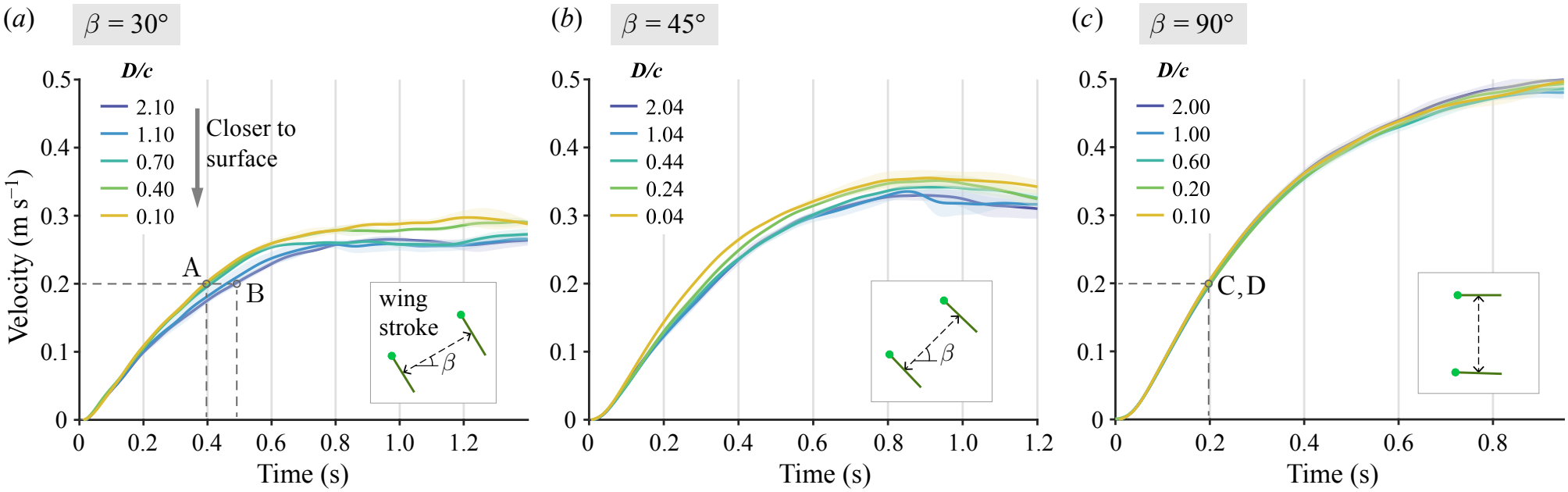
Effect of wing-surface distance on straight-line robot motion. Velocity responses at stroke plane angles (*a*) *β* = 30°, (*b*) *β* = 45°, and (*c*) *β* = 90°for varying normalized wing-surface distances *D*/*c*. We report mean velocity (*n* = 3) for each trial (solid curve) and one standard deviation (shaded region). At shallow stroke-plane angles (*β* = 30°, *β* = 45°), decreasing *D*/*c* increases acceleration and terminal velocity, with minimal effect at *β* = 90°. Points A-D mark near-surface (points A, C) and far-surface (points B, D) conditions at a common forward velocity *v* = 0.2 m s^−1^, highlighting greater near-surface acceleration at *β* = 30° compared with the nearly unchanged response at *β* = 90°. High-speed videography of all trials are detailed in **electronic supplementary material, movie S4**.

Looking specifically at a constant forward velocity *v* = 0.2 m s^−1^, we see that stroke-plane inclination affects near-surface propulsion enhancement (points A–D in figure 5*a,c*). For *β* = 30°, the near-surface condition (*D*/*c* = 0.1, point A) demonstrated greater acceleration than the far surface condition (*D*/*c* = 2.1, point B). In contrast at *β* = 90°, points C and D show similar acceleration.

To assess whether changes in hydrodynamic resistance could explain the increased acceleration and higher terminal velocities, we compared side-view high-speed recordings at near- and far-surface conditions. We observed no measurable differences in meniscus depth, water-ripple patterns, or robot pose between *D*/*c* = 0.1 and *D*/*c* > 9, as shown in **electronic supplementary material, movie S3**. These observations suggest that hydrodynamic resistance remained approximately independent of *D*/*c* at a given *v*, indicating that improved locomotion results primarily from enhanced aerodynamic thrust.

### 3.2. Aerodynamic thrust enhancement

We focused on the ground-effect-induced change in horizontal thrust at *β* = 30°, where straight-line experiments showed the strongest response to wing-surface distance (figure 5*a*). At terminal velocity, aerodynamic thrust balances the resistance acting on the robot (equation (2.2)). We used the longitudinal model to estimate the effective aerodynamic thrust (*T*_eff_) at the corresponding terminal velocities measured in figure 5*a* and compared these estimates with the far-from-surface baselines (*T*_0_(*v*_ter_, *β* = 30°)) to quantify the contribution of ground effect.

To account for the velocity dependence, we calculated the difference between the estimated thrust (*T*_eff_(*v*_ter_, *β* = 30°, *D*/*c*))and the baseline thrust-velocity (*T*_0_(*v*_ter_, *β* = 30°)) at each wing-surface clearance (figure 6*a*). The results are non-monotonic, with mean thrust decreasing as *D*/*c* decreases from 4.1 to 1.1, and then rising sharply for *D*/*c* < 0.7, with the largest change in mean thrust occurring at the smallest clearance. Similar trends were observed in previous studies [15, 18] and attributed to strong downwash from the previous stroke, which can lower the wing’s effective angle of attack and thus the aerodynamic force. This comparison indicates that the far-from-surface thrust–velocity relationship alone cannot account for the observed terminal velocities, supporting an additional aerodynamic thrust contribution from ground effect. Direct wind-tunnel measurements provided an independent assessment of this enhancement (figure 6*b*). The near-surface condition (*D*/*c* = 0.2) generated greater mean thrust than the far-surface condition (*D*/*c* = 9) across the tested freestream velocities. Mean thrust decreased with increasing freestream velocity under both conditions, but the relative enhancement varied with velocity. At *v* = 0.2 m s^−1^, reducing the wing–surface distance from *D*/*c* = 9 to 0.2 increased mean thrust by approximately 17%. Unlike the model-based estimates obtained at different terminal velocities, these direct measurements and interpolations quantify the effect of ground proximity at matched forward velocities.

**Figure 6.**
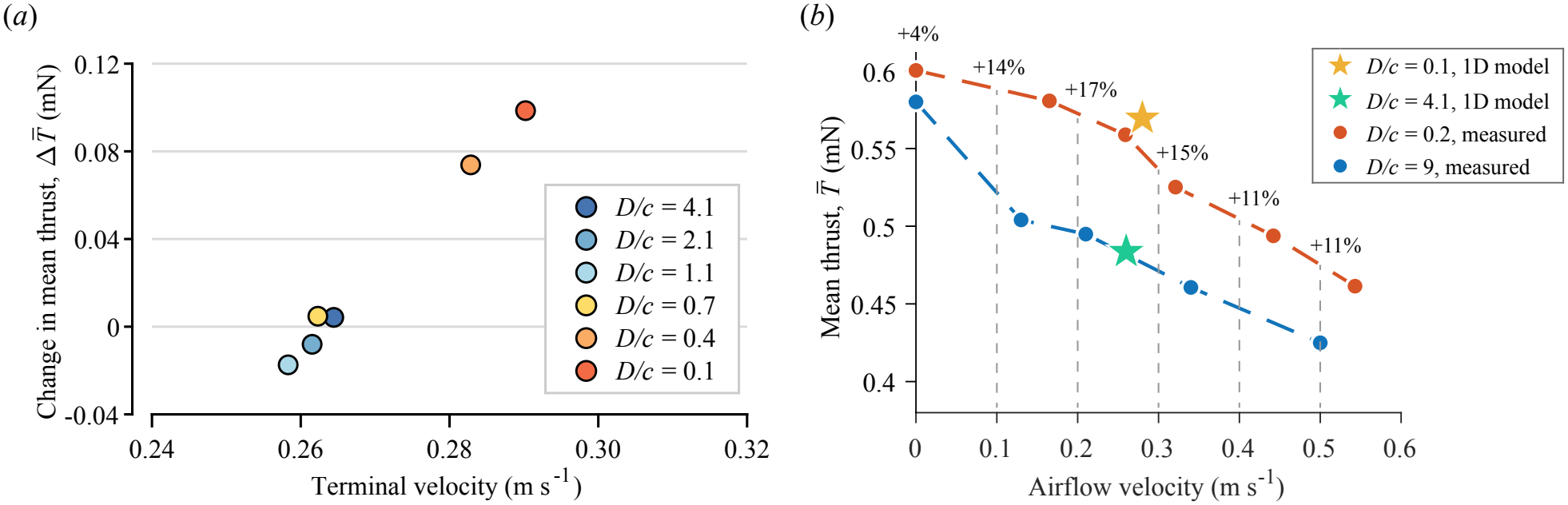
Observed changes in mean thrust with varying wing-surface distances at fixed *β* = 30°. (*a*) Estimated thrust enhancement at various wing-surface distances and corresponding terminal velocities (*v*_ter_). The change in mean thrust is the difference between the estimated *T*_eff_(*v*_ter_, *β* = 30*°, D*/*c*) and the baseline thrust-velocity relationship *T*_0_(*v*, *β* = 30*°*) obtained from direct force measurements. These results indicate a proximity-dependent thrust enhancement, with the largest differences between estimated mean thrust and the baseline at *D*/*c* = 0.4, 0.1. (*b*) Direct measurement of aerodynamic thrust enhancement at *β* = 30°. The measured mean thrust as a function of airflow velocity for near-surface (*D*/*c* = 0.2) and far-from-surface (*D*/*c* = 9) conditions is shown in red and blue, respectively (mean computed over two trials, each trial with ~ 130 flapping cycles). Dashed lines indicate linear interpolation. Star markers show the model-estimated effective thrust at the terminal velocities measured during straight-line locomotion at *D*/*c* = 0.1 (yellow) and *D*/*c* = 4.1 (green). The model estimates lie near the measured thrust for near- and far-from-surface conditions. Percentage annotations indicate the relative thrust enhancement at matched freestream velocities at 0.1 m s^−1^ intervals.

We also compared the model-estimated thrust at terminal velocity with the direct wind-tunnel measurements (star markers in figure 6*b*). The estimates at *D*/*c* = 0.1 and 4.1 lie near the corresponding near- and far-surface thrust–velocity curves at their respective terminal velocities. Although the wing–surface clearances differ between the two experimental setups, this comparison supports the consistency of the locomotion-based estimates with the directly measured aerodynamic forces.

### 3.3. Thrust estimation from momentum balance

The direct wind-tunnel measurements establish that ground proximity enhances horizontal thrust at *β* = 30°. To assess whether the measured flow fields reflect this enhancement and examine its dependence on stroke-plane angle, we calculated mean horizontal thrust per unit span from the two-dimensional PIV velocity fields using a momentum balance [40]. We compared near-surface (*D*/*c* = 0.2) and far-from-surface (*D*/*c* = 9) conditions at *β* = 30° and *β* = 90°, with a fixed freestream velocity of 0.2 m s^−1^.

We defined a fixed rectangular control area *A* enclosing the wing-swept region with boundary *L* (figure 7*a*). Although the momentum balance approach is theoretically independent of the size and placement of the control area [41, 42], in practice, experimental estimates are constrained by PIV resolution. We tested three boundary placements, *L*_I_–*L*_III_, where the upstream and upper boundaries of *L*_I_ extend by > 1*c*; of *L*_II_ by > 1*c* and 0.5*c*, respectively; and of *L*_III_ by 0.5*c* and > 1*c*, respectively. For all conditions, the downstream boundary extends by more than 2*c*, and the lower boundary is positioned above the moving surface. We selected the downstream boundary to minimize measurement errors associated with regions of high velocity gradients and concentrated vorticity [43, 44]. To maintain sufficient spatial resolution of the near-wing flow, we used a relatively small PIV field of view, which constrained the available control-area dimensions. We positioned the downstream boundary at the farthest available location, beyond the primary wake structures (figure 9). Assuming incompressible flow, negligible wing thickness, and neglecting out-of-plane momentum transport, we express the cycle-averaged force per unit span as

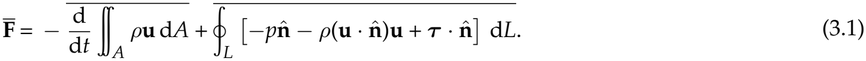

where **u** = (*u, v*, 0) is the in-plane velocity, *p* is pressure, *µ* is the dynamic viscosity of air, 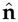 is the outward unit normal vector on the four edges of *L*, and ***τ*** = *µ*(***∇*u** + (***∇*u**)^*⊤*^) is the viscous stress tensor. The overline indicates cycle-averaged forces. We calculate the momentum flux for each PIV frame before averaging over the wingbeat cycle. We estimate the mean thrust 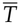 from the horizontal component of 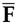.

**Figure 7.**
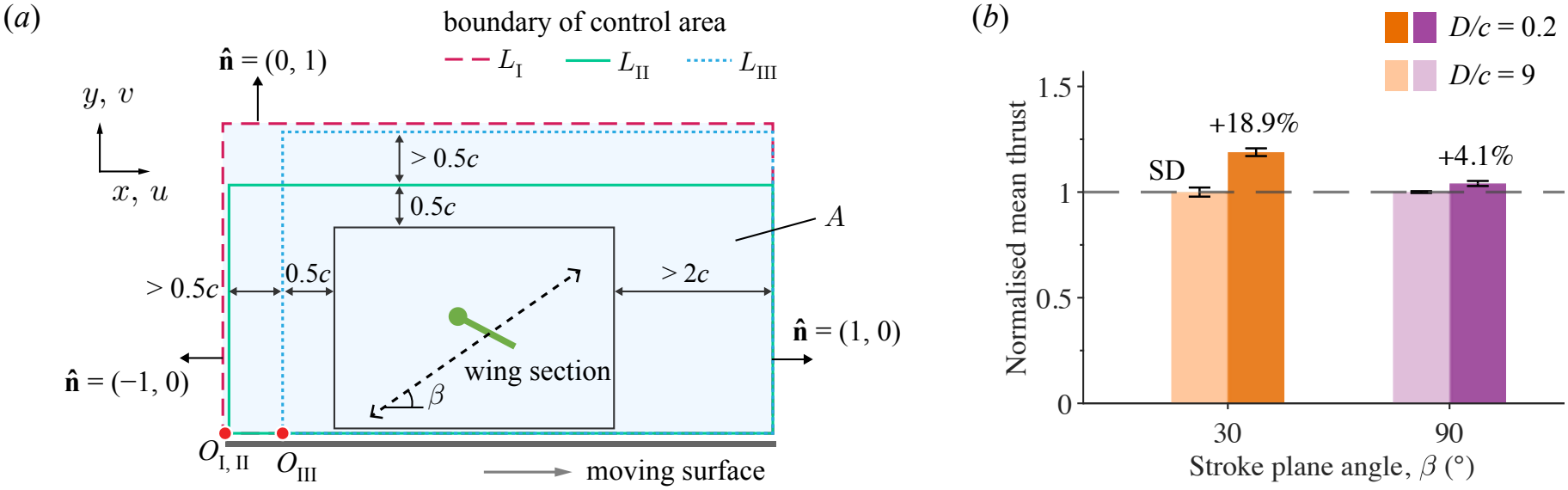
PIV-based estimates of mean horizontal thrust. (*a*) Control areas (*A*) used to estimate mean horizontal thrust per unit span from two-dimensional velocity fields. Each control area encloses the wing-swept region (black box), with boundaries positioned relative to the wing chord *c*. We compared three control areas with boundaries *L*_I_–*L*_III_. The unit normals 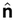 of all boundaries align with the two axes, and the position vector ***X*** is measured from the origin ***O*** located at the bottom left corner of the control area. (*b*) Estimated mean horizontal thrust per unit span at *β* = 30°and 90°, normalised by the corresponding far-from-surface baseline (*D*/*c* = 9). Reducing wing-surface clearance to *D*/*c* = 0.2 increases estimated thrust by an average of 18.9% at *β* = 30°, compared with 4.1% at *β* = 90°across the three control areas. Error bars indicate one standard deviation. All measurements were obtained at a freestream velocity of 0.2 m s^−1^.

**Figure 8.**
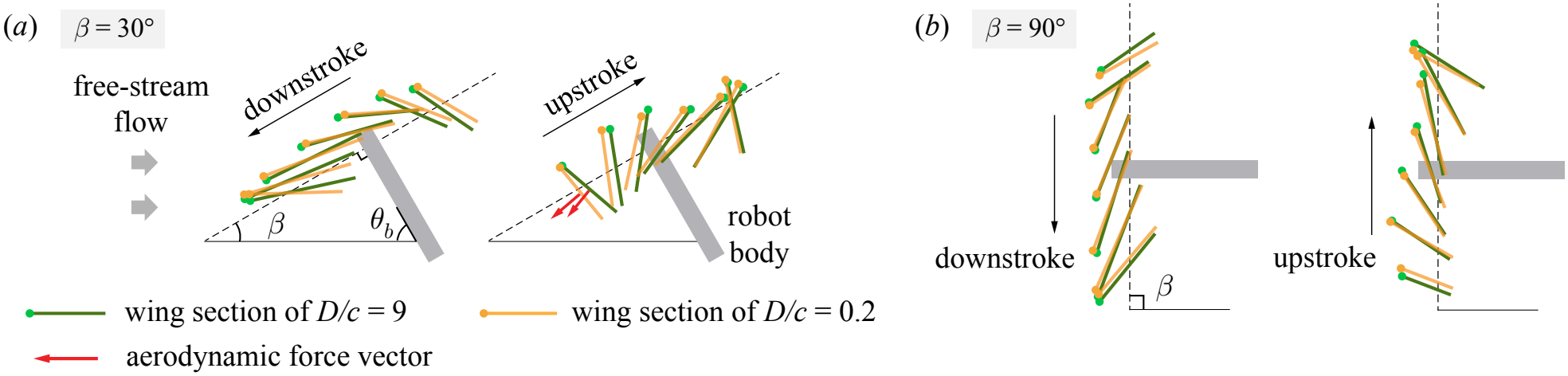
Effect of wing-surface proximity on sectional wing kinematics at (*a*) *β* = 30°and (*b*) *β* = 90°. We reconstructed the trajectories at near-surface (*D*/*c* = 0.2) and far-from-surface (*D*/*c* = 9) conditions by plotting every fifth frame during one wingbeat. (*a*) At *β* = 30°, the near-surface wing section deviates from the far-surface trajectory near the end of the downstroke, displaying advanced pitching. After stroke reversal, the near-surface wing exhibits delayed motion and altered orientation. (*b*) The trajectories nearly coincide at *β* = 90°. Arrows in (*a*) illustrate the assumed aerodynamic force vector at the indicated phase, suggesting a greater forward component in the near-surface condition. We observed little variations in flapping motion between cycles within each condition.

**Figure 9.**
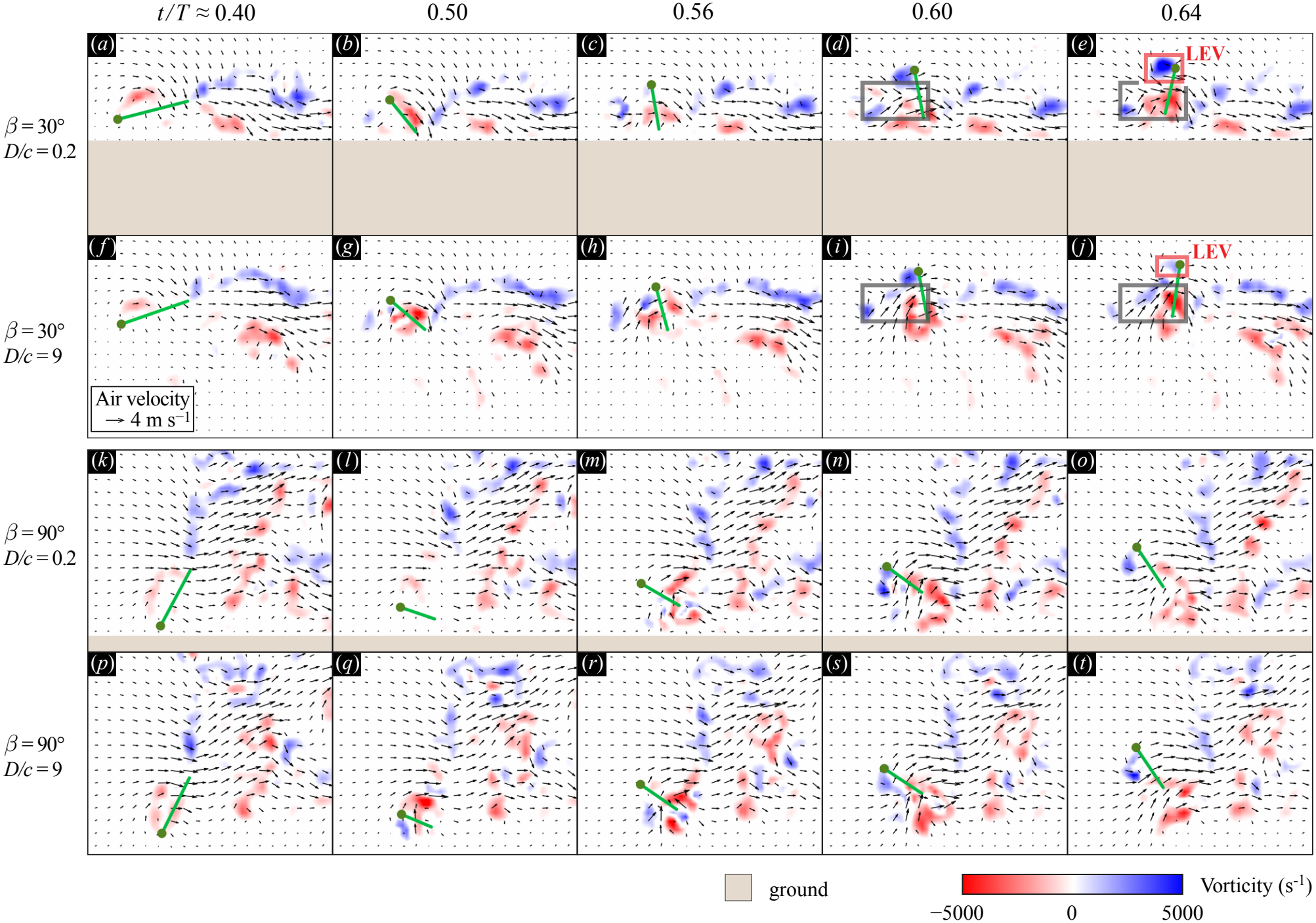
Effect of wing-surface proximity on near-wing flow during stroke reversal. Phase-resolved vorticity and velocity fields at *r* = 0.6*R* for (*a*–*e*) *β* = 30°, *D*/*c* = 0.2; (*f* –*j*) *β* = 30°, *D*/*c* = 9; (*k* –*o*) *β* = 90°, *D*/*c* = 0.2; and (*p*–*t*) *β* = 90°, *D*/*c* = 9. From left to right, the panels represent time sequence *t* /*T ≈* 0.40, 0.50, 0.56, 0.60, and 0.64. These phases capture the fluid field evolution around stroke reversal near *t* /*T* = 0.50, where 0 *≤ t* /*T* < 1 is the normalized timing in flapping period *T*. Colours indicate vorticity (blue: anticlockwise rotation; red: clockwise rotation), and superimposed vectors indicate the local airflow velocity. Green lines indicate tracked wing sections, with dots marking the leading edge. At *β* = 30°, the near-surface condition exhibits a stronger backwards-velocity component after stroke reversal compared to the far-surface condition (grey boxes in (*d* –*e*) and (*i*–*j*)). We also observe earlier leading-edge vortex (LEV) development during the initial upstroke for the near-surface case (red boxes in (*e*) and (*j*)). At *β* = 90°, the flow fields show little change with wing-surface proximity. All measurements use a freestream velocity of 0.2 m s^−1^. Please see PIV videos for *β* = 30°(**electronic supplementary material, movie S5**) and for *β* = 90°(**electronic supplementary material, movie S6**) for further details.

Because the PIV measurements do not directly provide pressure, we use the vortex-based identity from Noca et al.[41] to evaluate the pressure contribution:

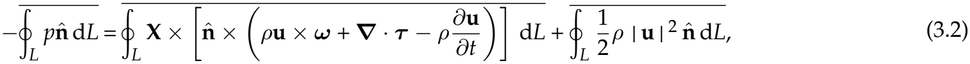

where **X** = (*x, y*, 0) is the position vector and ***ω*** = ***∇*** *×* **u** is the vorticity. We averaged the momentum balance over 6 wingbeats for each control area to reduce cycle-to-cycle variations. The approximately periodic wing motion and flow fields suggest that the cycle-averaged contributions from the unsteady terms (time derivatives in equation (3.1) and equation (3.2)) are small.

Using control area *A*_I_, the estimated mean horizontal thrust per unit span (mean *±* SD, *n* = 6 wingbeats) was 26.66 *±* 5.00 and 31.44 *±* 6.03 mN m^−1^ for the far- and near-surface conditions at *β* = 30°, and 56.72 *±* 5.18 and 59.07 *±* 2.11 mN m^−1^ at *β* = 90°. We test how the choice of control area affects the thrust estimate using control areas *A*_II_ and *A*_III_, which shift the upstream and upper boundaries, respectively, closer to the wing-swept region. The data across three control areas shows good consistency, as shown in (figure 7*b*). At *β* = 30°, reducing *D*/*c* from 9 to 0.2 increased the estimated mean thrust per unit span by 18.9% *±* 1.0%, compared with only 4.1% *±* 1.0% at *β* = 90°. The small variation across control areas demonstrates that the estimated enhancement is relatively insensitive to the tested boundary placements.

Although the absolute thrust per unit span was greater at *β* = 90°, the relative enhancement due to ground proximity was substantially larger at *β* = 30°. This contrast supports the stronger sensitivity of horizontal thrust to ground proximity at shallow stroke-plane inclinations, as seen in the straight-line locomotion performance (§3.1). Furthermore, the ~19% enhancement is comparable to the measured 17% increase with the same nominal conditions (§3.2). Although the two-dimensional approximation neglects out-of-plane momentum transport, the the comparable enhancements provide quantitative support for the relationship between near-wing flow modification and increased horizontal thrust at shallow stroke-plane angles.

### 3.4. Wing kinematics and near-wing flow

We observed that the greater relative thrust enhancement at shallow stroke-plane angles is accompanied by changes in wing motion and surrounding flow. We compared sectional wing kinematics at *β* = 30° and 90° under near-surface *D*/*c* = 0.2 and far-from-surface *D*/*c* = 9 conditions, with a freestream velocity of 0.2 m s^−1^ (figure 8*a,b*). At *β* = 30°, the near- and far-surface wing sections differed near wing reversal. Near the end of the downstroke, the leading-edge positions nearly coincided, but the near-surface wing section rotated into its upstroke orientation earlier. This advanced pitch rotation may promote interactions with the wake generated during the preceding downstroke [45] and influence leading-edge circulation development [22]. Immediately after stroke reversal, the near-surface wing section exhibited a more vertical orientation, increasing the horizontal component of the aerodynamic force, which acts normal to the wing [46]. The subsequent lag in near-surface wing motion before the trajectories converged suggests a change in aerodynamic loading on the passively rotating wing. In contrast, the wing-section trajectories at *β* = 90° remained nearly symmetric throughout the flapping cycle (figure 8*b*).

To examine how changes in wing motion relate to the surrounding flow, we compared the phase-resolved velocity and vorticity fields near the downstroke-to-upstroke reversal (figure 9). At *β* = 30°, the near-surface condition produced a larger backward velocity component in front of the wing immediately after stroke reversal, as highlighted in figure 9*d–e,i–j*, and detailed in **electronic supplementary material, movie S5**. Here, we define “backward” as the direction of relative airflow, opposite to robot’s forward direction. The near-surface flow also exhibited earlier leading-edge vortex (LEV) development during the initial upstroke (figure 9*e,j*). This stronger LEV is also consistent with the motion lag during initial upstroke we observed in figure 8*a*, as the LEV corresponds to low-pressure region on wing leeward surface. In contrast, the velocity and vorticity fields at *β* = 90° remained qualitatively similar between the two wing-surface distances (figure 9*k–o,p–t*, also in **electronic supplementary material, movie S6**). These observations suggest that ground proximity modifies the flow generated during wing reversal, potentially influencing aerodynamic force production.

The cycle-averaged wake [17] provides a broader view of how the stroke-plane angle affects flow behavior (figure 10*a– d*). At *β* = 30°, the far-surface wake descended and spread vertically downstream (figure 10*a*), whereas the near-surface wake remained concentrated above the boundary, with stronger backward velocity (figure 10*b*). At *β* = 90°, the near- and far-surface wakes showed comparatively little change in profile and distribution (figure 10*c,d*). These contrasting responses indicate that ground proximity modifies both wing kinematics and wake development more strongly at shallow stroke-plane angles, consistent with the larger relative thrust enhancement measured at *β* = 30°.

**Figure 10.**
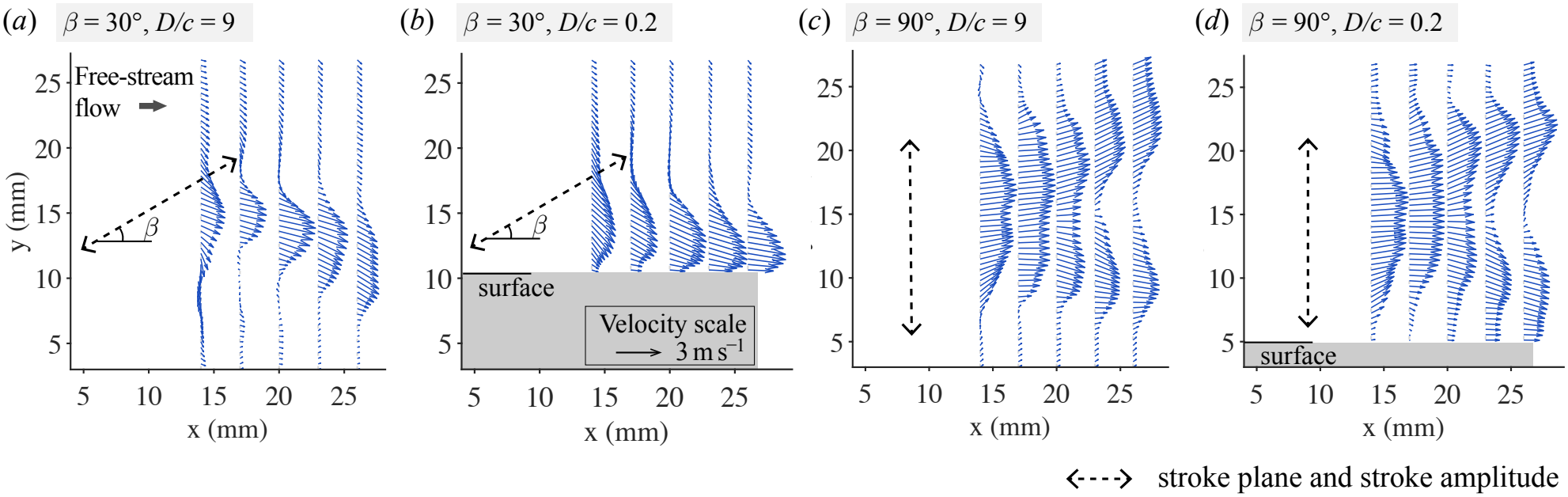
Effect of wing-surface proximity on cycle-averaged wake. (*a-d*) Cycle-averaged wake velocity fields (blue vectors) for (*a*) *β* = 30°, *D*/*c* = 9; (*b*) *β* = 30°, *D*/*c* = 0.2; (*c*) *β* = 90°, *D*/*c* = 9; and (*d*) *β* = 90°, *D*/*c* = 0.2. Dashed double-headed arrows indicate stroke-plane inclination and stroke amplitude of the leading edge motion (they do not reflect the actual wing-surface distance). At *β* = 30°, ground proximity concentrates the backward-directed wake near the boundary, whereas the wake shows minimal change at *β* = 90°.

## 4. Discussion

### 4.1. Wing-surface aerodynamic interactions

Our results demonstrate that the aerodynamic benefits of near-surface flapping depend strongly on stroke-plane inclination, in addition to wing-surface proximity. At shallow inclinations, the wing can approach the boundary over a longer portion of the stroke, potentially strengthening interactions between the wing-generated flow and the nearby surface. The passive wing pitch angle of our robotic platform allows wing motion to respond to aerodynamic loading, rather than prescribing wing kinematics independently of the surrounding flow. Our results show that the aerodynamic response to ground proximity involves changes in both wing motion and surrounding flow, and we leave the analysis of their individual contribution to thrust enhancement to future work.

The stronger backward flow and earlier LEV development observed after stroke reversal provide additional insight into the aerodynamic interactions. These features are qualitatively similar with flow generated during the fling phase of the “clap-and-fling” mechanism [47], where wing separation promotes LEV development and enhances force production [26]. However, the interaction between a flapping wing and a nearby rigid boundary differs from that between two moving wings. Whether the observed flow changes arise primarily from altered wing kinematics, direct wing-boundary vortex interactions, or their combined effects remains an open question.

Future experiments could distinguish these effects by comparing passive and prescribed wing rotation under otherwise identical operating conditions. Independently controlling wing pitch would allow us to evaluate how much of the thrust enhancement persists when wing stroke remains unchanged, while varying wing and hinge stiffness could reveal how structural compliance affects the response. Combining these experiments with time-resolved force measurements and three-dimensional flow visualization would help connect the observed changes in wing motion and vortex development to aerodynamic force generation throughout the flapping cycle.

### 4.2. Implications for interfacial flight

Our findings raise the possibility that aquatic insects could modulate aerodynamic ground effect through changes in body posture or wing kinematics. Whether waterlily beetles actively exploit these interactions during interfacial flight, particularly when transitioning between interfacial and airborne locomotion, remains an open question.

The robotic platform reproduces several important operating conditions of waterlily beetle interfacial flight (table 1). The robot operates at a stroke-plane inclination of *β* = 30°, within the 20°–35° range observed in waterlily beetles, and at comparable normalized wing–surface clearance (*D*/*c≈* 0.2). Its forward velocity (0.33 m s^−1^) and Strouhal number (2.9) are also comparable to those reported for beetles (0.30 m s^−1^ and 2.3, respectively). However, the robot has larger wings, a smaller stroke amplitude, and a higher Reynolds number (*Re* = 1230 versus 480). These differences limit direct quantitative extrapolation of the measured thrust enhancement to living insects, but the comparable operating conditions support the robot’s use as a physical model for studying near-surface flapping aerodynamics.

**Table 1.**
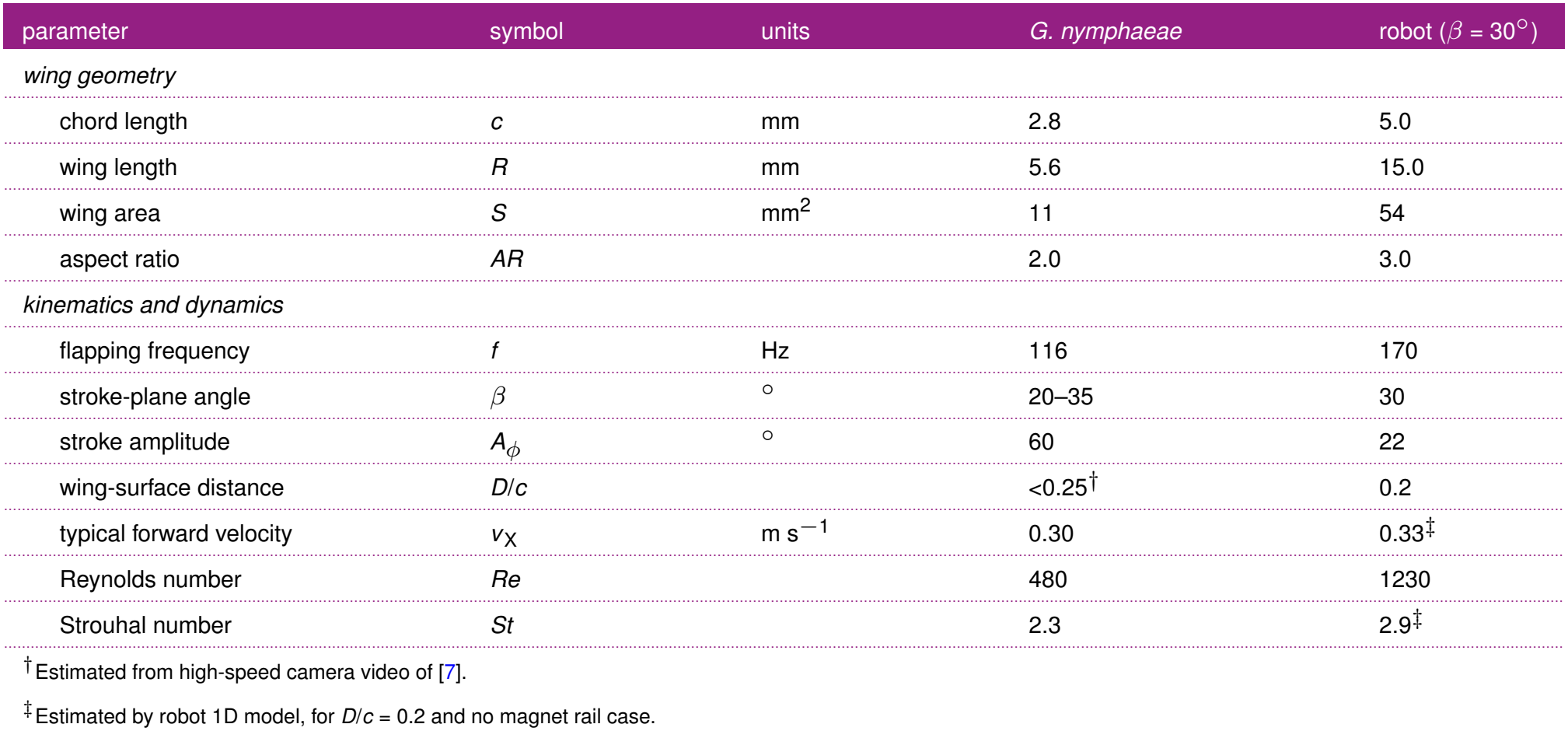
Comparison of waterlily beetle species (*G. nymphaeae* [7]) and the flapping-wing robot.

| parameter | symbol | units | *G. nymphaeae* | robot (*β* = 30°) |
|---|---|---|---|---|
| *wing geometry* |  |  |  |  |
| chord length | *c* | mm | 2.8 | 5.0 |
| wing length | *R* | mm | 5.6 | 15.0 |
| wing area | *S* | mm<sup>2</sup> | 11 | 54 |
| aspect ratio | *AR* |  | 2.0 | 3.0 |
| *kinematics and dynamics* |  |  |  |  |
| flapping frequency | *f* | Hz | 116 | 170 |
| stroke-plane angle | *β* | ° | 20–35 | 30 |
| stroke amplitude | *A*<sub>*ϕ*</sub> | ° | 60 | 22 |
| wing-surface distance | *D*/*c* |  | <0.25<sup>†</sup> | 0.2 |
| typical forward velocity | *v*<sub>X</sub> | m s<sup>−1</sup> | 0.30 | 0.33<sup>‡</sup> |
| Reynolds number | *Re* |  | 480 | 1230 |
| Strouhal number | *St* |  | 2.3 | 2.9<sup>‡</sup> |

<sup>†</sup>Estimated from high-speed camera video of [7].
<sup>‡</sup>Estimated by robot 1D model, for *D*/*c* = 0.2 and no magnet rail case.

An important distinction between our experiments and biological interfacial flight is the nature of the underlying boundary. We used rigid surfaces to isolate the aerodynamic effects of wing–surface proximity, whereas aquatic insects flap above a deformable air–water interface. Surface deformation, wave generation, and differences in fluid boundary conditions may modify the interaction between wing-generated flow and the interface. Comparing flapping aerodynamics above rigid and deformable boundaries remains an area for future experiments.

### 4.3. Implications for robot design

Wing-surface proximity is a potential design parameter to enhance horizontal propulsion in insect-scale flapping-wing robots. By adjusting stroke-plane inclination and wing-surface clearance, robots may exploit aerodynamic interactions with nearby boundaries without increasing actuation amplitude or frequency. This could enable lower actuation voltages for a given thrust requirement or improve acceleration during intermittent flapping strategies [48], in which active propulsion alternates with passive gliding or rest on the water surface. Whether these benefits translate into reduced electrical power consumption or improved cost of transport remains to be determined.

However, operating close to a boundary introduces constraints on wing motion and robot stability [49]. Maintaining the desired clearance requires precise control of body posture and height, particularly in environments with surface disturbances. For interfacial robots, excessive body oscillations or surface waves could also cause wing contact with water, reducing performance or damaging onboard electronics. Future work should therefore examine how robots can regulate stroke-plane inclination and wing–surface clearance to balance aerodynamic thrust enhancement with stability, control requirements, and energy consumption.

### 4.4. Concluding remarks

Our experiments demonstrate that near-surface flapping can enhance horizontal thrust in insect-scale flapping-wing systems, with the magnitude of enhancement strongly dependent on stroke-plane inclination. By combining straight-line locomotion, longitudinal dynamic modelling, direct force measurements, and PIV, we establish a quantitative connection between ground proximity, aerodynamic thrust, and robot motion. The accompanying changes in sectional wing kinematics and near-wing flow provide insight into the aerodynamic interactions associated with this enhancement. More broadly, this work demonstrates the value of insect-scale robotic physical models for studying aerodynamic interactions that are difficult to isolate in living organisms.

Several questions remain regarding the underlying force-generation mechanisms. Our two-dimensional PIV measurements do not capture spanwise flow or resolve instantaneous aerodynamic forces, limiting our ability to determine the contributions of altered wing motion and vortex development. Future studies combining three-dimensional flow measurements, time-resolved force measurements, and independently controlled wing kinematics could distinguish these effects and establish predictive models of ground effect across wing geometries, stroke-plane inclinations, and operating conditions.

## Ethics

The work did not require approval from human subjects or animal-welfare ethics committee.

## Data Accessibility

All data necessary to review the manuscript can be found at Zenodo [50].

## Declaration of AI Use

AI tools were used for grammar and readability, as well as assistance with data-analysis code implementation using methods conceptualized by authors. The authors manually checked all code before use and take full responsibility for the results.

## Authors’ Contributions

CZ: conceptualization, data curation, formal analysis, investigation, methodology, validation, visualization, writing - original draft; CU: data curation, methodology; HG: data curation, methodology; ZJW: conceptualization, investigation, methodology, supervision, validation, writing - review and editing; EFH: conceptualization, data curation, funding acquisition, investigation, methodology, project administration, supervision, writing - review and editing.

## Acknowledgements

We thank Sean Donlon for customizing the glass plates, Prof. Kevin Chen for helpful discussion on PIV measurements. We also thank Dr. Jiefu Yin, Cindy Martin, and Stephanie Pierson for coordinating the laser safety setup for PIV experiments.

