## Supplementary Material for "Near-surface flight enhances thrust in an insect-inspired flapping-wing robot"

### Cubic approximation of thrust as a function of forward speed

Chenyu Zhang

November 2026

Using a two-dimensional (2D) wing model and a quasi-steady estimate [1], we show that the cycle-averaged thrust of an inclined flapping wing can be approximated by a third-order polynomial in the body forward speed $u$, when $u$ is small compared with the wing sweep speed $w$ relative to the body. We consider the far-from-surface condition and neglect ground effect.

#### Effective angle of attack

As shown in figure S1, the stroke plane is inclined at a fixed angle $0° \le \beta \le 90°$ from the horizontal. The plane's normal points upward at $\beta = 0°$ (as in normal hovering) and forward at $\beta = 90°$. The geometric wing pitch $\alpha$ is the angle between the wing chord and the stroke direction. For simplicity, we set a constant wing sweep speed $w = U_{\mathrm{rms}}$, and use the same fixed pitch $\alpha$ on both half-strokes. The body moves through still air at speed $u$. Both $w$ and $u$ denote non-negative magnitudes.

**Downstroke.** Applying the law of cosines to the velocity triangle in figure S1(a) gives the effective incoming speed during downstroke:

$$v_{\mathrm{eff,dn}} = \sqrt{w^2 + u^2 + 2uw\cos\beta}. \tag{S1}$$

Forward motion also changes the effective angle of attack. Since $\alpha_{\mathrm{eff,dn}} = \alpha - \delta_{\mathrm{dn}}$, we have

$$\begin{aligned}\sin\alpha_{\mathrm{eff,dn}} &= \sin\alpha\cos\delta_{\mathrm{dn}} - \cos\alpha\sin\delta_{\mathrm{dn}} = \frac{w\sin\alpha + u\sin(\alpha-\beta)}{v_{\mathrm{eff,dn}}},\\ \cos\alpha_{\mathrm{eff,dn}} &= \cos\alpha\cos\delta_{\mathrm{dn}} + \sin\alpha\sin\delta_{\mathrm{dn}} = \frac{w\cos\alpha + u\cos(\alpha-\beta)}{v_{\mathrm{eff,dn}}}.\end{aligned} \tag{S2}$$

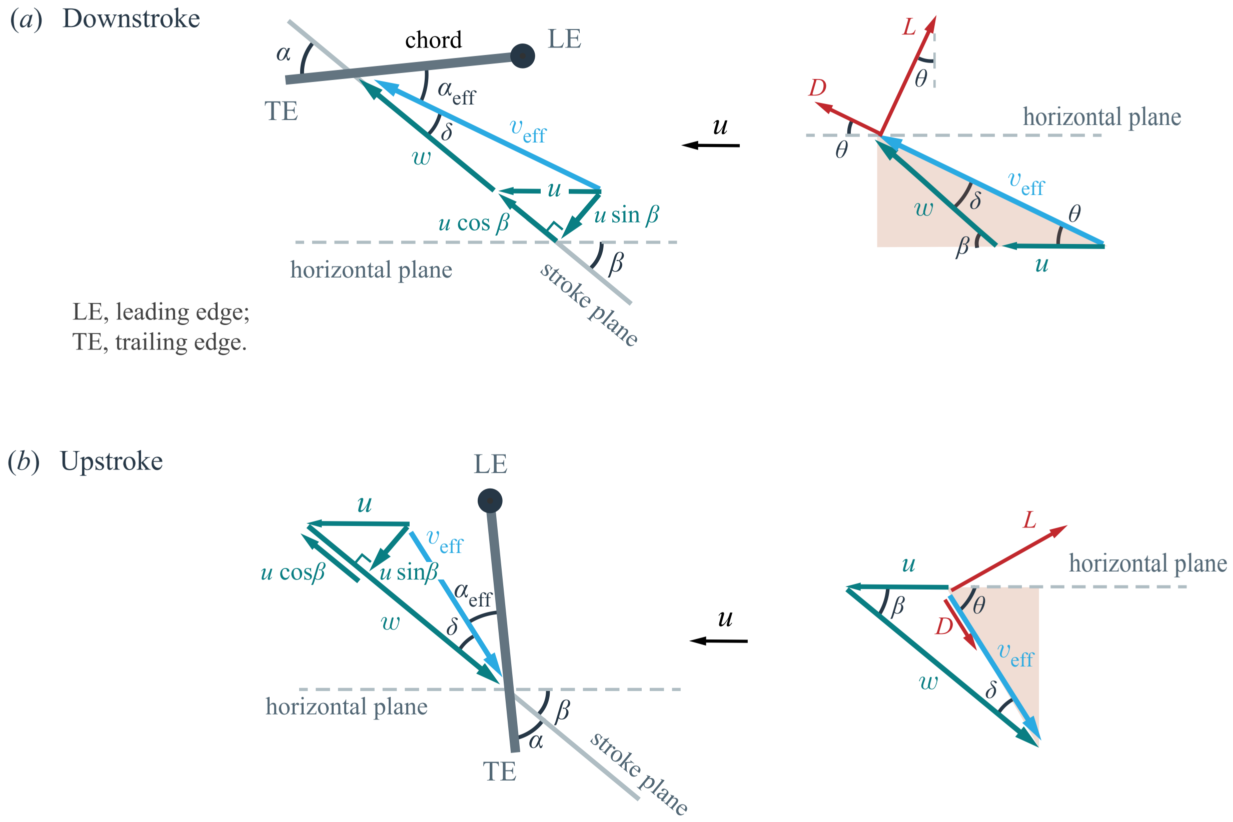


Figure S1: Stroke geometry, effective angle of attack, and aerodynamic forces on a 2D chord during ($a$) downstroke and ($b$) upstroke. and the tangent to its stroke direction. $\alpha$, geometric wing pitch. $\beta$, stroke plane angle. $u$, flow relative to wing due to body motion. $w$, flow relative to wing due to wing motion. $v_{\text{eff}}$ , effective incoming flow speed. $\alpha_{\text{eff}}$ , effective angle of attack. $\delta$, the angle between the incident flow and the stroke direction. $\theta$, the angle between the incident flow and the horizontal. Both wing lift $L$ and wing drag $D$ contribute to the thrust (the net horizontal force).

**Upstroke.** Similarly, the geometry in figure S1(b) gives

$$\begin{aligned} v_{\text{eff,up}} &= \sqrt{w^2 + u^2 - 2uw\cos\beta}, \\ \sin\alpha_{\text{eff,up}} &= \frac{w\sin\alpha - u\sin(\alpha+\beta)}{v_{\text{eff,up}}}, \\ \cos\alpha_{\text{eff,up}} &= \frac{w\cos\alpha - u\cos(\alpha+\beta)}{v_{\text{eff,up}}}. \end{aligned} \tag{S3}$$

At $u = 0$, both speeds reduce to $w$, and both effective angles reduce to the geometric pitch $\alpha$.

#### Forward thrust

We use the following quasi-steady lift and drag coefficients for a 2D wing [1]:

$$C_L = 1.2\sin(2\alpha_{\text{eff}}), \quad C_D = 1.4 - \cos(2\alpha_{\text{eff}}). \tag{S4}$$

The effective angle of attack generally differs between upstroke and downstroke. The aerodynamic coefficients for downstroke $C_{L,\text{dn}}, C_{D,\text{dn}}$ and upstroke $C_{L,\text{up}}, C_{D,\text{up}}$ are obtained by substituting equations (S2) (S3) into equation (S4), respectively.

For air density $\rho$ and chord length $c$, the downstroke lift and drag per unit span are

$$L_{\text{dn}} = \frac{1}{2}\rho\, c\, v_{\text{eff,dn}}^2 C_{L,\text{dn}}, \qquad D_{\text{dn}} = \frac{1}{2}\rho\, c\, v_{\text{eff,dn}}^2 C_{D,\text{dn}}. \tag{S5}$$

Projecting both forces onto the forward direction gives the downstroke thrust

$$\begin{aligned} T_{\text{dn}} &= L_{\text{dn}}\sin\theta_{\text{dn}} - D_{\text{dn}}\cos\theta_{\text{dn}} \\ &= \frac{1}{2}\rho c\, v_{\text{eff,dn}}\big[C_{L,\text{dn}} w\sin\beta - C_{D,\text{dn}}(u + w\cos\beta)\big], \end{aligned} \tag{S6}$$

where $\sin\theta_{\text{dn}}$ and $\cos\theta_{\text{dn}}$ are given by the coloured triangle in figure S1(a).

Similarly, the thrust during upstroke is

$$T_{\text{up}} = \frac{1}{2}\rho c\, v_{\text{eff,up}}\big[C_{L,\text{up}} w\sin\beta + C_{D,\text{up}}(w\cos\beta - u)\big]. \tag{S7}$$

Assuming the two half-strokes have equal duration, the cycle-averaged thrust is

$$\overline{T} = \frac{1}{2}T_{\text{dn}} + \frac{1}{2}T_{\text{up}}. \tag{S8}$$

#### Thrust in terms of the advance ratio

We define the advance ratio as $J = u/w$, the ratio of body forward speed to flapping speed [2]. Dividing the incoming flow speeds by $w$ gives

$$\frac{v_{\text{eff,dn}}}{w} = \sqrt{1 + 2J\cos\beta + J^2}, \qquad \frac{v_{\text{eff,up}}}{w} = \sqrt{1 - 2J\cos\beta + J^2}. \tag{S9}$$

Equations (S6) and (S7) then take the following form, where $f_{\mathrm{dn}}(J), f_{\mathrm{up}}(J)$ are quadratic polynomials in $J$ with coefficients determined by $\alpha$ and $\beta$:

$$\begin{aligned}\frac{T_{\mathrm{dn}}}{\frac{1}{2}\rho c w^2} &= \frac{1}{\sqrt{1+2J\cos\beta+J^2}}\; f_{\mathrm{dn}}(J),\\ \frac{T_{\mathrm{up}}}{\frac{1}{2}\rho c w^2} &= \frac{1}{\sqrt{1-2J\cos\beta+J^2}}\; f_{\mathrm{up}}(J).\end{aligned} \tag{S10}$$

#### Cubic approximation

We use $(1+x)^{-1/2} = 1 - \frac{1}{2}x + \frac{3}{8}x^2 - \frac{5}{16}x^3 + O(x^4)$ to expand the factors in equation (S10):

$$\frac{1}{\sqrt{1\pm 2J\cos\beta+J^2}} = 1 \mp J\cos\beta + \frac{3\cos^2\beta-1}{2}J^2 \pm \frac{3\cos\beta-5\cos^3\beta}{2}J^3 + O(J^4). \tag{S11}$$

For our robot, $|J| < 1/3$. Over this range, the expansion converges, and truncating after the cubic term gives an error below $< 1\%$. The fourth term's $\left|J^4\right| \sim 0.01$ is substantially small. Combining equations (S8)–(S11) and keeping terms up to the cubic $J^3$ gives

$$\boxed{\begin{aligned}\frac{\overline{T}}{\frac{1}{2}\rho c w^2} \approx\;& 1.2\sin\beta\sin 2\alpha\\ &+ \left[\left(3.4\cos^2\beta - 1.4\right)\cos 2\alpha - 1.4\left(1+\cos^2\beta\right)\right]\cdot J\\ &+ \left(0.2 + 1.8\cos^2\beta\right)\sin\beta\sin 2\alpha\cdot J^2\\ &- \left(0.3\cos 2\alpha + 0.7\right)\sin^4\beta\cdot J^3, \qquad J = \frac{u}{w}.\end{aligned}} \tag{S12}$$

The geometric wing pitch $\alpha$ enters equation (S12) through $\sin(2\alpha)$ and $\cos(2\alpha)$, a consequence of the terms in $C_L$ and $C_D$ (equation (S4)). Thus, $\overline{T}$ is approximately a cubic polynomial in $u$.

**Check: Zero forward speed.** At $u = 0$, the cycle-average thrust reduce to

$$\frac{\overline{T}}{\frac{1}{2}\rho c w^2} \approx (1.2\sin\beta\sin 2\alpha) \;\propto\; \sin\beta. \tag{S13}$$

The proportional dependence on $\sin\beta$ is expected, as the wing drag cancels over a period, and the force is normal to the stroke plane [1]. The mean aerodynamic force is perpendicular to the stroke plane. To compare with experiments, we use measured values from figure S2 at $u = 0$, and use the single point of $\beta = 90°$ to determine the sine prediction (grey curve in figure S3). The curve agrees well with the other four measurements.

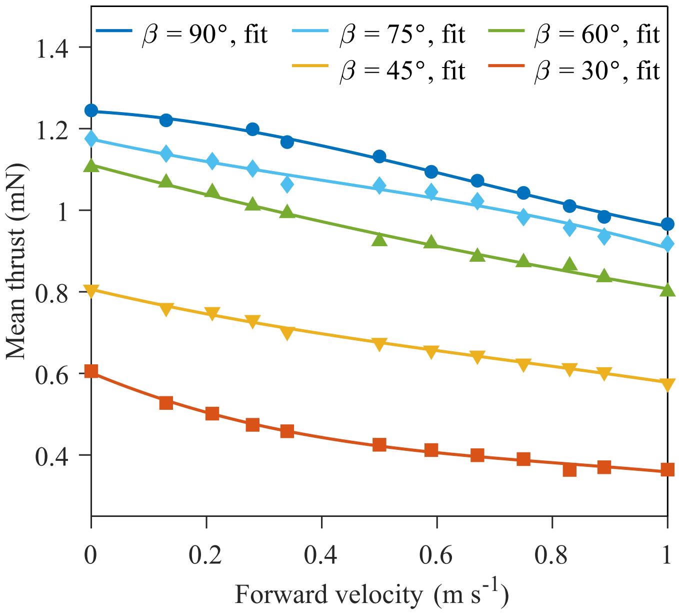


Figure S2: Measured mean thrust as a function of body forward velocity at five stroke-plane angles $\beta$. Each point represents the mean thrust of measurements over $\sim$ 260 flapping cycles. The curves are third-order polynomial fits.

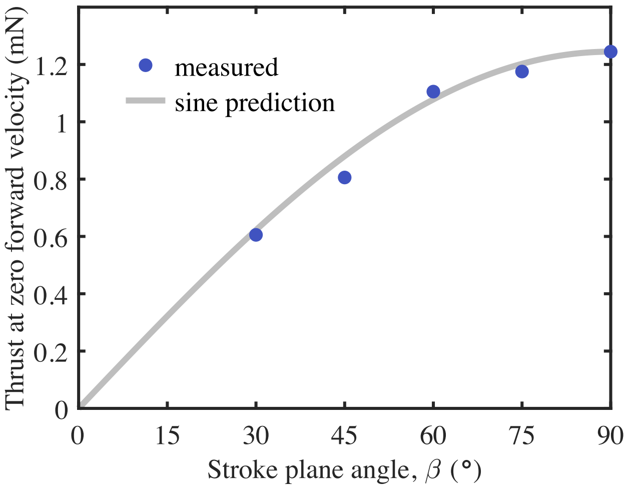


Figure S3: Measured and predicted thrust at zero forward speed as a function of $\beta$. The points show the measurements from figure S2 at $u = 0$, and the sine curve is determined by the measurement at $\beta = 90°$. The predicted curve agrees well with the remaining four points.

#### Representative stroke inclinations

**Horizontal stroke ($\beta = 0°$).** It is the same stroke geometry as normal hovering (recovers the hovering when $J = 0$). Lift has no horizontal component. The net thrust comes entirely from the difference in wing drag between the two half-strokes, and is linear to $J$:

$$\frac{\overline{T}}{\frac{1}{2}\rho c w^2} = -2\,(1.4 - \cos(2\alpha))\,J = -2\,C_D\,J < 0. \tag{S14}$$

The "negative thrust" indicates that the force opposes forward motion. Equation (S14) gives the same result as [3] equation (4.1).

**Vertical stroke ($\beta = 90°$).** The two half-strokes now have the same effective speed, $w\sqrt{1+J^2}$. Equation (S12) reduces to

$$\frac{\overline{T}}{\frac{1}{2}\rho c w^2} \approx 1.2\sin(2\alpha) - 1.4[1+\cos(2\alpha)]J + 0.2\sin(2\alpha)J^2 - [0.7 + 0.3\cos(2\alpha)]J^3. \tag{S15}$$

**Inclined stroke ($\beta = 30°$).** For the shallowest stroke inclination tested, equation (S12) gives

$$\frac{\overline{T}}{\frac{1}{2}\rho c w^2} \approx 0.6\sin(2\alpha) + [1.15\cos(2\alpha) - 2.45]J + 0.775\sin(2\alpha)J^2 - \frac{7+3\cos(2\alpha)}{160}J^3. \tag{S16}$$

The cubic coefficient is small at $\beta = 30°$, as $\sin^4\beta$ in equation (S12) drops rapidly with decreasing $\beta$.

For stroke inclination $30° \leq \beta \leq 90°$ and fixed $0° < \alpha < 90°$, the third order prediction shows that mean horizontal thrust decreases monotonically over $0 < J \leq 1/3$, which aligns with the experiments in figure S2. These results motivate our choice of a third-order polynomial, with the coefficients fitted to the measured mean thrust over the experimental speed range (figure S2).
